# Human immunodeficiency virus type 2 capsid protein mutagenesis reveals amino acid residues important for virus particle assembly

**DOI:** 10.1101/2022.01.31.478542

**Authors:** Huixin Yang, Nathaniel Talledge, William G. Arndt, Wei Zhang, Louis M. Mansky

**Affiliations:** Institute for Molecular Virology, University of Minnesota – Twin Cities, Minneapolis, MN 55455 USA; Division of Basic Sciences, School of Dentistry, University of Minnesota – Twin Cities, Minneapolis, MN 55455 USA; Masonic Cancer Center, University of Minnesota – Twin Cities, Minneapolis, MN 55455 USA; Comparative Molecular Biosciences Graduate Program, University of Minnesota – Twin Cities, St. Paul, MN 55108, USA; Biochemistry, Molecular Biology & Biophysics Graduate Program, University of Minnesota – Twin Cities, Minneapolis, MN 55455 USA; Characterization Facility, College of Sciences and Engineering, University of Minnesota – Twin Cities, Minneapolis, MN 55455 USA

**Keywords:** retrovirus, lentivirus, virus-like particle, maturation, morphology

## Abstract

Human immunodeficiency virus (HIV) Gag drives virus particle assembly. The capsid (CA) domain is critical for Gag multimerization mediated by protein-protein interactions. The Gag protein interaction network defines critical aspects of the retroviral lifecycle at steps such as particle assembly and maturation. Previous studies have demonstrated that the immature particle morphology of HIV-2 is intriguingly distinct relative to that of HIV-1. Based upon this observation, we sought to determine the amino acid residues important for virus assembly that might help explain the differences between HIV-1 and HIV-2. To do this, we conducted site-directed mutagenesis of targeted locations in the HIV-2 CA domain of Gag and analyzed various aspects of virus particle assembly. A panel of 31 site-directed mutants of residues that reside at the HIV-2 CA inter-hexamer interface, intra-hexamer interface and CA inter-domain linker were created and analyzed for their effects on the efficiency of particle production, particle morphology, particle infectivity, Gag subcellular distribution and *in vitro* protein assembly. Seven conserved residues between HIV-1 and HIV-2 (L19, A41, I152, K153, K157, N194, D196) and two non-conserved residues (G38, N127) were found to significantly impact Gag multimerization and particle assembly. Taken together, these observations complement structural analyses of immature HIV-2 particle morphology and Gag lattice organization as well as provide important comparative insights into the key amino acid residues that can help explain the observed differences between HIV immature particle morphology and its association with virus replication and particle infectivity.

## Introduction

Human immunodeficiency virus (HIV) belongs to the lentivirus genus of the *Retroviridae*. To date, HIV has infected more than 70 million people worldwide [1]. HIV comprises two main types, HIV type 1 (HIV-1) and HIV type 2 (HIV-2), which clearly have similar modes of replication, transmission, and clinical symptomatology [2]. Despite these similarities, there are notable differences in the general prevalence, rate of transmission, and relative particle infectivity between these viruses. For example, HIV-2 has less widespread and is mainly found in West Central Africa, with reduced infectivity and transmissibility [2, 3]. HIV-2 is also known to have lower viral loads during the course of the infection, slower rates of CD4^+^ T-cell decline, and delayed clinical progression to AIDS [4]. Individuals infected with HIV-2 are typically less infectious in the early stages of infection and can more readily lead to long-term nonprogressors [5, 6]. Different steps in the HIV life cycle have been successfully targeted by combined antiretroviral therapy (ART), leading to a remarkable reduction in overall morbidity and mortality [7].

The HIV assembly pathway represents a promising and generally underdeveloped target for antiretroviral intervention. While intensive efforts have been made to target HIV assembly [8–12], approved antiretroviral drugs to date have been elusive. The HIV capsid (CA) has been viewed as a particularly promising antiviral drug target due to the fact that its stability and integrity are critical for virus replication and particle infectivity [13]. For example, lenacapavir (LEN, GS-6207, GS-CA2), an investigational first-in-class long-acting HIV-1 CA inhibitor, is currently in phase 2/3 clinical trials and is seeking regulatory approval [14]. LEN binds to the hydrophobic pocket formed by two adjoining CA subunits within the CA hexamer. Since HIV-2 is not susceptible to many anti-HIV-1 drugs [15, 16], insights into the molecular basis for antiretroviral drug susceptibility can inform the development of effective, next-generation treatments for both HIV types.

The HIV Gag polyprotein is the major structural protein responsible for driving retrovirus assembly [12]. The HIV-1 (NL4-3) and HIV-2 (ROD) Gag proteins have a moderate level of amino acid sequence conservation (i.e., 59% identity, 72% similarity). HIV assembly engages three key Gag domains: the matrix (MA) domain is responsible for Gag-membrane binding [17, 18]; the nucleocapsid (NC) domain interacts with the viral RNA packaging signal and encapsidates the viral genome [19]; and the CA domain is critical for Gag multimerization during virus assembly and core formation during virus maturation. The CA domain has two structurally distinct subdomains, *i.e.*, the CA amino-terminal subdomain (CANTD) and the CA carboxy-terminal subdomain (CACTD) [20], which are connected by a flexible inter-domain linker. The major homology region (MHR), a 20-residue segment (amino acid residues 153-172 in HIV-1 CA) toward the amino-terminus of the CACTD, is highly conserved among retroviruses. During particle assembly, Gag (Pr55^Gag^) oligomerizes at the plasma membrane, forming a radially arranged lattice that curves and deforms the plasma membrane [17]. Formation of the immature Gag lattice drives budding and release of immature particles. The virally-encoded protease (PR) cleaves the Gag polyprotein to create MA, CA and NC proteins that are needed to form a mature virion [21]. The MA protein remains associated with the inner leaflet of the viral membrane. The CA protein reassembles into a cone-shaped core that contains a dimeric viral genomic RNA coated with NC protein [17].

The morphologies of HIV-1 immature and mature particles have been intensively studied [22, 23]. Previous observations indicate that the mature HIV-1 capsid is made from a lattice of ∼250 CA hexamers closed by the insertion of 12 CA pentamers into a fullerene cone structure [24]. In the immature HIV-1 Gag lattice, no pentamers have been found; the Gag hexamers are interspersed with gaps to accommodate the curvature of the viral membrane. The CA hexamer of the immature HIV-1 Gag lattice contains an inner ring of six CACTD subdomains held together by contacts between adjacent CA molecules [25, 26]. The interactions between CA monomers are related by six-fold symmetry within a hexamer (i.e., intra-hexamer) (**Fig. S1A**) and by two-fold (**Fig. S1B, S1C**) and three-fold symmetry (**Fig. S1D**) between neighboring hexamers (i.e., inter-hexamer). The hexamers are stabilized by interactions at these interfaces [24].

Many immature Gag lattice structures have been published to date, including Rous sarcoma virus (RSV) [27], Murine leukemia virus (MLV) [28], and HIV-1 [29]. It is well recognized that key amino acid residues in CA can dictate virus particle morphology. Mutagenesis studies have been used to confirm the importance of the interface residues for core morphology and stability and virion infectivity [24, 25, 30, 31]. Mutations in the CA domain of Gag have been shown to affect particle assembly and virion infectivity in different retroviruses, including HIV-1 [32], Simian immunodeficiency virus (SIV) [33], Mason-Pfizer monkey virus (M-PMV) [34], and RSV [35]. Several key mutants, for example the HIV-1 CA W184A and M185A cause Gag assembly defects, produce non-infectious virus particles, reduce CA dimerization and intermolecular Gag–Gag binding constants *in vitro*, diminish immature particle production *in vivo* [30, 36]; similar studies have been done with SIV [33]. In addition to introduction of disruptive site-directed mutants, interchange of residues between closely related viruses have been used to assess effects of the mutations on particle production and infectivity. For example, a previous study of HIV-1 and SIVmac residues in HIV-1 CA Helix 7, such as E128N, I141C, were found to cause severe reductions in particle production and infectivity when mutated to the corresponding SIVmac residue [37]. This study indicated that these non-conserved amino acids in HIV-1 CA Helix 7 have important roles in assembly and particle infectivity.

Given these observations, we hypothesized that the differences between HIV-1 and HIV-2 CA interactions could help explain the distinct morphological differences observed, as well as to provide insights into the role of particle assembly and morphology. To investigate this, as well as to establish residues important for virus assembly, a panel of site-directed mutants was analyzed for their impact on the efficiency of immature particle production. Selected mutants were additionally analyzed for mature particle production, infectivity, Gag subcellular distribution and *in vitro* protein assembly. These integrative analyses revealed several key amino acid residues and implicated novel HIV-2 CA interactions that are required for efficient virus particle assembly. Taken together, our studies provide molecular detail regarding key CA residues that mediate virus particle assembly, highlight fundamental differences between HIV-1 and HIV-2 CA, and provide further insights into fundamental aspects of HIV particle assembly that can inform intervention therapies.

## Results

### Identification of conserved HIV-2 CA residues that impact immature particle production

Previous studies on HIV-1, MLV, and RSV have indicated that specific amino acid residues in the Gag CA domain encode the key determinants that impact immature and mature virus particle morphology as well as implicate key CA-CA interactions [27-29, 38, 39]. To date, limited data regarding HIV-2 Gag CA residues involved in particle assembly have been described. We sought out to identify critical residues in HIV-2 Gag CA that mediate HIV particle assembly [30, 36, 40] (**Fig. 1**). We conducted alanine-scanning mutagenesis of 21 HIV-2 residues that were predicted to impact virus particle assembly based upon comparative analysis or based on structural predictions. We used immunoblot analysis in conjunction with a tractable HIV-2-like particle system to analyzed authentic, immature virus particle assembly and release (**Fig. 2**).

**Figure 1.**
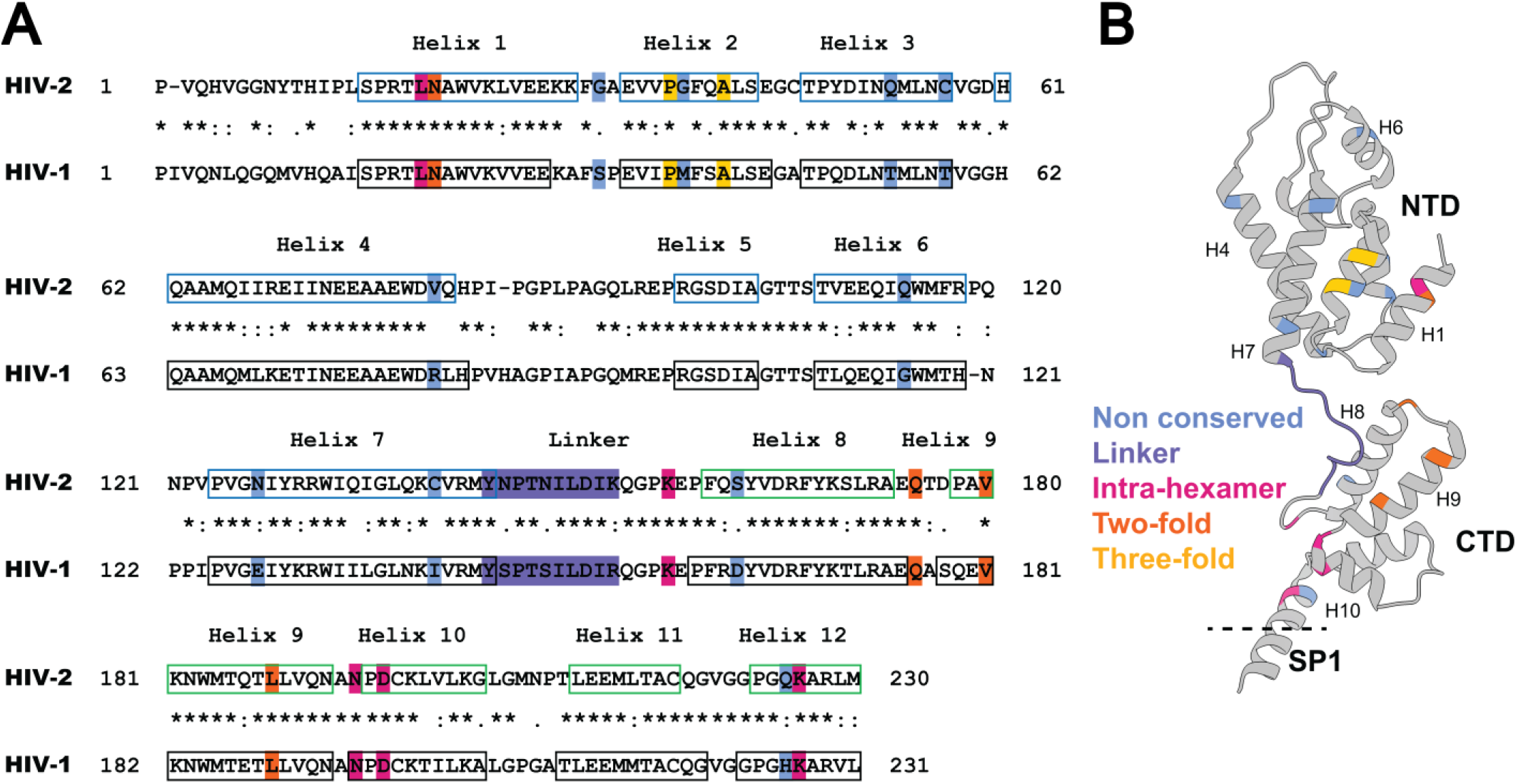
Sequence alignment and CA structure. (A) Alignment of the HIV-1 and HIV-2 CA sequences. Amino acid sequences in CA of HIV-1 NL4-3 (GenBank accession no. AF324493; CA 1-231; Gag 133-363) and HIV-2 ROD (GenBank accession no. M15390; CA 1-230; Gag 136-365) were aligned by using the Clustal Omega Multiple Sequence Alignment [57]. Locations of the HIV-1 immature CA helices in the alignment are based on previous studies [22] (PDB ID: 5L93) and are indicated by black boxes. The HIV-2 immature CANTD (CA 1-144; Gag 136-279) helices were previously described [58] (PDB ID: 2WLV) and are indicated with blue boxes. The CACTD (CA 145-230; Gag 280-365) helices are predicted by the PSIPRED server [59, 60], and are indicated by green boxes. “*” indicates conserved amino acids; “:” indicates amino acid substitution with high amino acid similarity; “.” indicates amino acid substitutions with low similarity. The blue, purple, magenta, orange, and gold-shaded amino acid residues highlight mutagenesis at non-conserved residues, linker, intra-hexamer interface, two-fold interface, and three-fold interface, respectively. (B) Presumably the ribbon diagram is of the CA monomer structure. Shown are CA amino-terminal domain (NTD), the carboxy-terminal domain (CTD), the spacer peptide 1 (SP1), and alpha-helices 1, 4, 6, 7, 8, 9, and 10 (i.e., H1, H4, H6, H7, H8, H9, and H10). The non-conserved, linker, intra-hexamer, two-fold and three-fold residues analyzed are indicated on the ribbon structure.

**Figure 2.**
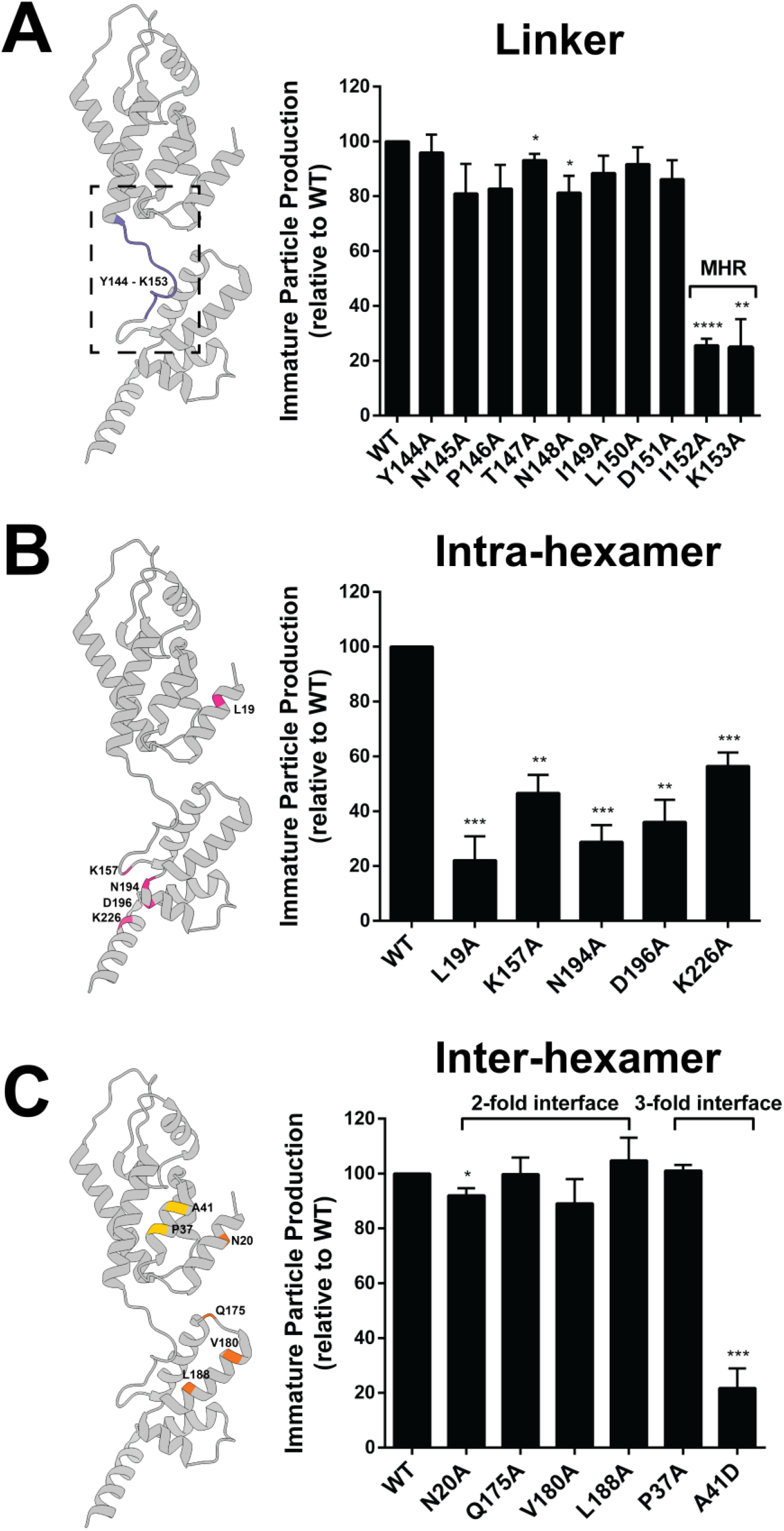
Immature particle production of HIV-2 CA mutants at the CANTD-CACTD linker (A), the intra-hexamer interface (B), and the inter-hexamer interface (C). 293T/17 cells were transfected with Plasmids expressing the HIV-2 Gag (WT or alanine-scanning mutants) were transfected into 293T/17 cells, and the cell culture supernatants were harvested 48h post transfection. Immunoblot analysis was conducted to determine the amount of immature particle production for the CA conserved mutants relative to that of WT. Histograms show the relative immature particle production for each HIV-2 CA mutant compared to WT (set to 100). Error bars represent standard error of the mean from three independent experiments. Significance relative to WT was determined by unpaired t test. ****, *P* < 0.0001; ***, *P* < 0.001; **, *P* < 0.01; *, *P* < 0.05. MHR = Major homology region. Mutation locations are colored in the HIV CA monomer structure, using the same color-coding scheme as shown in Figure 1B. Representative immunoblot images were shown in Figure S2.

Ten site-directed mutations were located at the HIV-2 CANTD-CACTD linker domain (**Fig. 2A**). Among these mutations, I152A and K153A were discovered to reduce immature particle production relative to WT by 3.9-fold, and 4-fold, respectively (**Fig. 2A**). These residues were located in the MHR region. Notably, some conserved residues in HIV-1 MHR (e.g., HIV-1 Q155N) led to a dramatic reduction in HIV-1 immature particle production [41]. However, the phenotype of mutant in HIV-1 CA (I153A) which corresponds to HIV-2 I152A has not been previously reported.

Five site-directed mutants of residues at the Gag intra-hexamer interface were analyzed. Four of these mutants (i.e., L19A, K157A, N194A, and D196A) resulted in a 2-fold or greater reduction in immature particle production relative to that of WT (**Fig. 2B**). The HIV-2 CA L19A and N194A mutants reduced particle production by 4.5-fold and 3.5-fold, respectively, compared to WT. The comparable residues have not been characterized to date in the HIV-1 CA domain. These mutations are novel, because the comparable residues in the HIV-1 CA domains have not been previously reported. The HIV-2 CA K157A and D196A mutants resulted in 2.1-fold and 2.8-fold reductions in particle production compared to WT, respectively. This observation is similar to that reported with comparable mutations in the HIV-1 CA domain on particle production [30, 36, 42].

Inter-hexamer interactions play a critical role in formation of the HIV-1 Gag lattice. Given this, four site-directed mutants at the putative HIV-2 two-fold inter-hexamer interface and two mutants at the three-fold inter-hexamer interface were analyzed to confirm their influence on HIV-2 particle production (**Fig. 2C**) based on CA structural analysis (**Fig. S1**). The two-fold inter-hexamer interface mutants had particle production efficiencies comparable to that of WT, with the exception for the N20A mutant, which had a minor reduction in particle production (∼92% of WT). One of the two mutants at the three-fold inter-hexamer interface, i.e., A41D, had a 4.6-fold reduction in particle production (**Fig. 2C**).

### Identification of non-conserved HIV-2 CA residues and their impact on immature particle production

Previous studies have demonstrated differences between HIV-1 and HIV-2 immature particles [39], suggesting a model in which relatively minor changes in protein-protein interaction geometry may profoundly impact overall particle morphology [39]. To further investigate this, a panel of 10 site-directed mutations at non-conserved CA residues were generated by mutating each HIV-2 CA residue to the corresponding HIV-1 amino acid residue (**Fig. 3A**).

**Figure 3.**
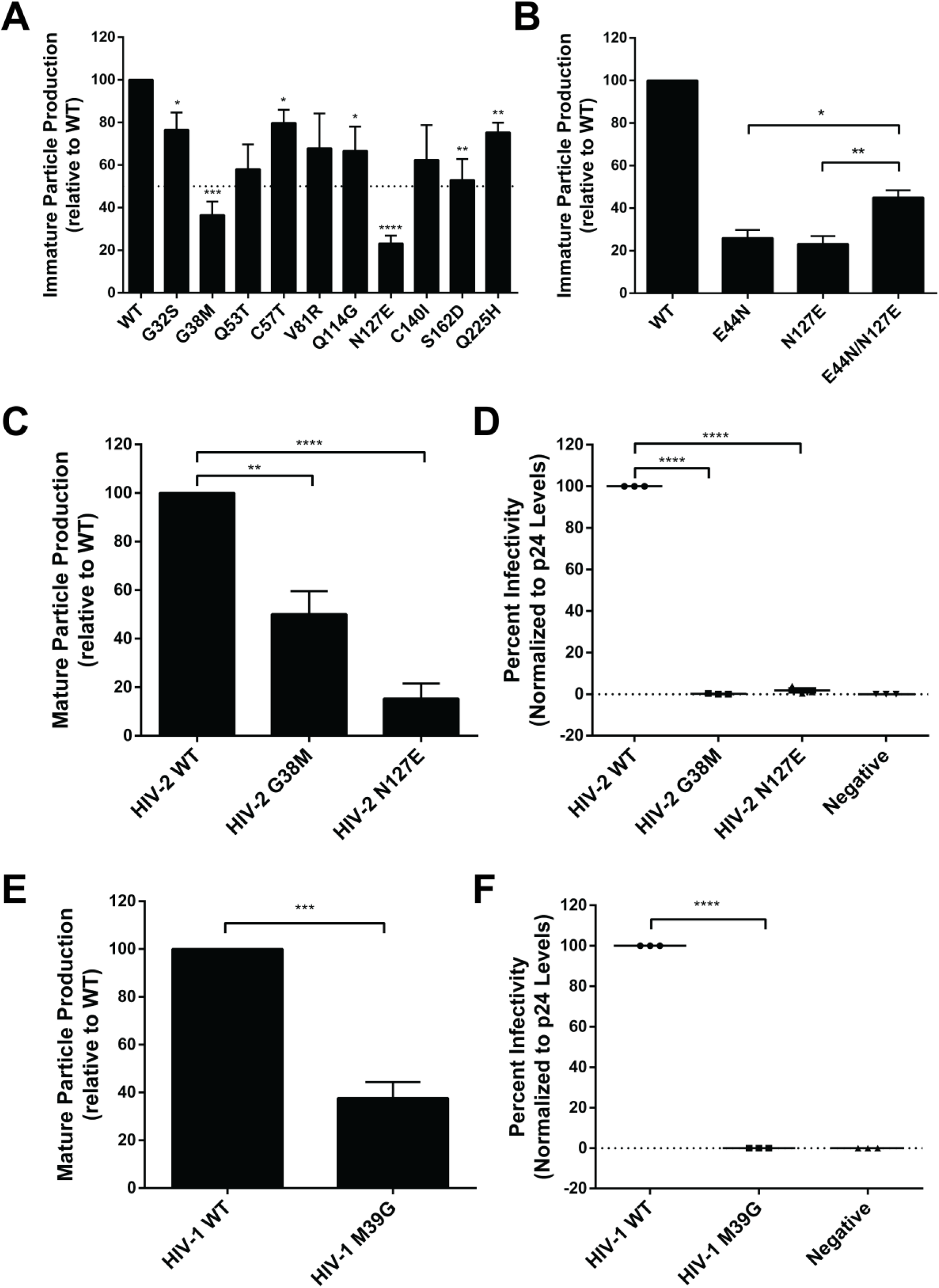
Particle production and infectivity analysis for HIV CA mutants at non-conserved amino acid residues. (A) Immature particle production of HIV-2 CA mutants at non-conserved amino acid residues. Particle production of mutants from a Gag expression construct is relative to that of WT (set at 100). (B) Immature particle production of HIV-2 CA mutants. Shown is particle production of the HIV-2 CA E44N, N127E and E44N/N127E mutants relative to that of WT (set at 100). (C) Mature HIV-2 particle production from a HIV-1 single-cycle vector for selected mutants. Shown is particle production of the HIV-2 G38M and N127E CA mutants relative to that of WT (set at 100). (D) Infectivity of HIV-2 CA mutants. Shown is the infectivity relative to that of WT (set at 100) for the HIV-2 G38M and N127E mutants. (E) Mature particle production from a HIV-1 single-cycle vector of a HIV-1 CA mutant. Shown is mature particle production for the HIV-1 M39G CA mutant relative to that of WT (set at 100). (F) Infectivity analysis of HIV-1 CA mutant. Immunoblot analysis of CA p24 levels was conducted to determine the amount of particle production for the M39G mutant relative to that of the WT HIV-1 single-cycle vector. Virus particle infectivity was determined by flow cytometry analysis of infected cells that express a fluorescence reporter gene from the integrated HIV-1 provirus. Mutant particle infectivity was relative to that of the WT HIV-1 vector virus and was normalized to CA p24 levels. Error bars represent standard deviations from three independent experiments. Significance relative to WT was determined by an unpaired t test. ****, *P* < 0.0001; ***, *P* < 0.001; **, *P* < 0.01; *, *P* < 0.05. Representative immunoblot images were shown in Figure S2.

Two HIV-2 CA mutants (i.e., G38M and N127E) reduced particle production by greater than 50% compared to that of WT. These amino acid residues are both located at the three-fold inter-hexamer interface in the CANTD, based upon structural comparison to the HIV-1 lattice structure. The HIV-2 G38M mutation reduced both immature particle production (**Fig. 3A**) and mature particle production (**Fig. 3C**). The equivalent residue in HIV-1 CA mutation (M39G) was also found to reduce particle production relative to that of WT (**Fig. 3E**).

It was previously observed that mutation of the HIV-1 CA E128 residue to the corresponding HIV-2 residue (i.e., HIV-1 E128N) reduced viral replication [37]. We observed the HIV-2 N127E mutant reduced immature particle production by 4.3-fold compared to WT (**Fig. 3A**), and resulted in a 6.7-fold reduction of mature particle production compared to WT (**Fig. 3C**). The HIV-1 E128N mutant led to a 2.6-fold reduction of mature particle production compared to WT. Analysis of the position of E128 in the HIV-1 immature CA cryo-EM structure [22] reveals that E128 is within interaction distance (<4 Å) with the E45 residue. Based upon the HIV-2 sequence, two single mutations (i.e., HIV-2 E44N, N127E) were generated to compare to that of WT (E44/N127). Production of immature particles for the E44N and N127E mutants were about 4-fold less than the WT (**Fig. 3B**). The HIV-2 E44N/N127E double mutant led to a doubling of immature particle production compared to the single mutants, but was 2-fold less than WT.

### Perturbation of Gag subcellular distribution

Gag subcellular distribution can alter Gag multimerization and productive particle assembly and release [17]. To determine if the subcellular distribution of HIV-2 Gag mutants was associated with reductions in immature particle production, selected mutants were engineered to contain an YFP tag at the carboxy-terminus of the Gag protein. The mutants analyzed included L19A (intra-hexamer interface), A42D (inter-hexamer interface), I152A (located at the CANTD-CACTD linker domain), and the non-conserved G38M and N127E mutants. HIV-2 WT and mutant Gag-YFP constructs were transfected into Hela cells, and analyzed for their ability to multimerize and form Gag puncta, particularly at the plasma membrane [39]. Among the five mutants analyzed, the HIV-2 G38M and N127E mutants led to diffuse fluorescence in the cytoplasm as well as localization in the nucleus (**Fig. 4**). These observations indicated that the G38M and N127E mutations (which are non-conserved residues in the three-fold interface) perturb Gag subcellular localization. Similar observations were also made in other retroviruses, including RSV [39, 43, 44] and walleye dermal sarcoma virus (WDSV) [39].

**Figure 4.**
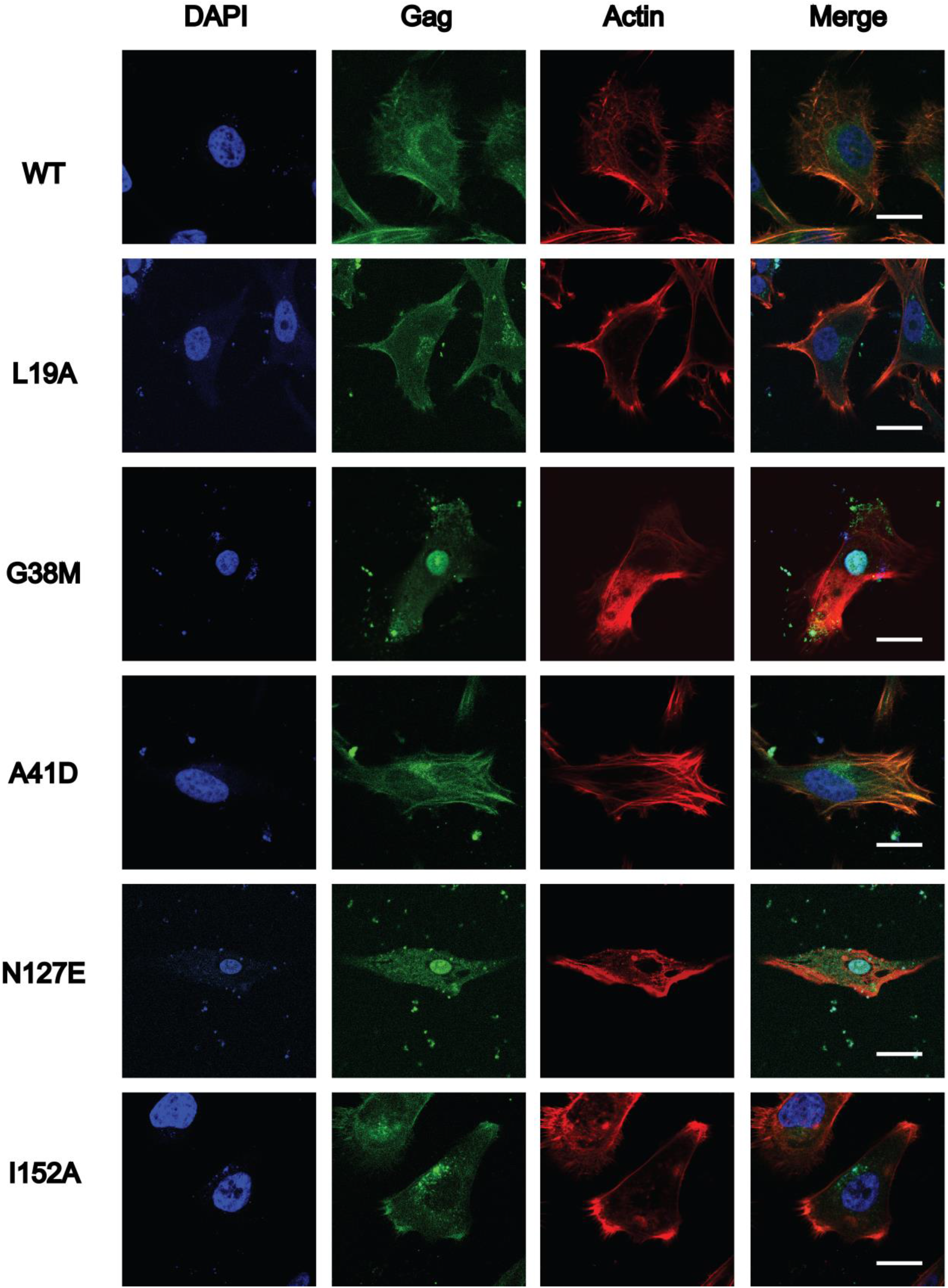
Subcellular localization of HIV-2 WT Gag and selected CA domain mutants. HeLa cells were transfected with either HIV-2 Gag-eYFP or a site-directed Gag CA domain mutant. Representative images for WT HIV-2 Gag-eYFP and each Gag CA domain mutant (*i.e*., L19A, G38M, A41D, N127E and I152A) are shown. At least 15 individual cells were imaged across three independent replicates for a total of 15 cells. Scale bar, 20 μm. Nuclei (blue; stained with DAPI), Gag (green) localization; actin (red, stained with actin red) are indicated, and the merged composite is shown.

### G38M and N127E alter HIV-2 immature particle morphology

Information regarding the critical residues of HIV-2 CA that dictate particle morphology is limited in comparison to that of HIV-1 CA. Previous reports have identified that the HIV-2 lattice appears more uniform with slightly larger particle diameters than that of HIV-1 particles or other retroviral particles [27–29, 38]. The striking morphological differences between immature WT HIV-1 and HIV-2 particles were notable and imply that relatively minor changes in the geometry of protein-protein interactions can significantly impact overall particle morphology [39].

The HIV-2 CA G38M and N127E mutants were next examined for their influence on immature particle morphology, and to contrast them to HIV-2 WT immature particles, which have a relatively uniform particle morphology [39] (**Fig. 5A-C**). In particular, HIV-2 WT immature particles have a tightly packed Gag lattice, implying a highly organized CA layer below the viral membrane. The majority of the HIV-2 WT immature particles were spherical, with a small population having other morphologies (**Fig. 5A**). In contrast, HIV-2 G38M (**Fig. 5B**) and N127E (**Fig. 5C**) immature particles were observed to have an increase in non-spherical morphologies with clear gaps and defects in the underlying Gag lattice. Additionally, the mean particle diameter was significantly larger for the two mutants than HIV-2 WT immature particles (**Fig. 5D**). These observations suggests that the G38M and N127E mutations impose a defect in Gag multimerization to form a proper Gag lattice structure.

**Figure 5.**
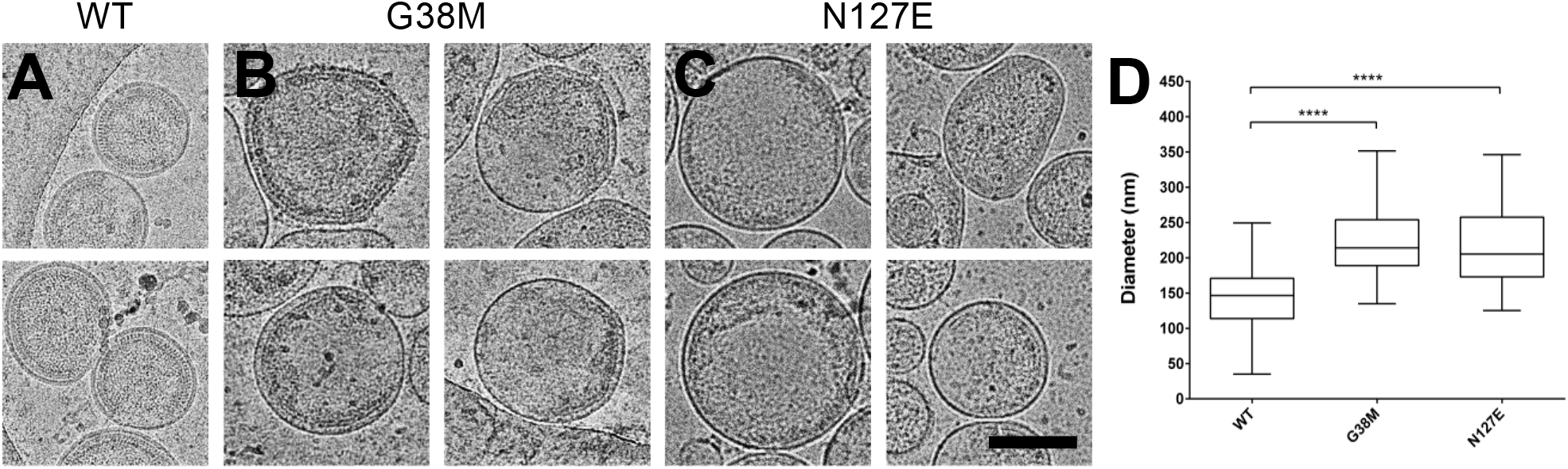
Cryo-electron microscopy of immature HIV-2 WT or Gag CA domain mutant particles. The HIV-2 Gag expression plasmid or selected mutant was transfected into 293T/17 cells, and the particles produced from the cells were concentrated and purified from cell culture supernatants prior to cryo-electron microscopy analysis. Representative images are shown for the (A) WT and the (B) G38M and (C) N127E CA domain mutants (scale bar, 150 nm). (D) Diameter quantification of the WT, G38M, and N127E immature particles (n > 50) generated by calculating the mean along two perpendicular axes per particle. The histogram shows the range, median, and the 25th to 75th percentiles of the particle diameters. Significance relative to WT was determined by an unpaired t test. ****, *P* < 0.0001.

### The G38M and N127E mutations severely diminish HIV-2 particle infectivity

Infectivity assays of CA point mutants help determine whether a single amino acid can alter virus replication. Here, a single-cycle infectivity system with a fluorescent reporter system was used as previously described [45]. Vector virus particles (WT or mutant) were harvested from producer cells and used to infect permissive target cells and assessed by flow cytometry. Relative infectivity of mutants compared to WT was determined.

The HIV-2 G38M and N127E mutants exhibited severe defects in particle infectivity, 100-fold and 56-fold reductions compared to WT, respectively (**Fig. 3D**). To assess if the equivalent mutations in HIV-1 (see **Fig. 1A**) impact particle infectivity, the site-directed mutant HIV-1 CA M39G was analyzed and was found to have a 100-fold reduction in particle infectivity compared to WT (**Fig. 3F**). A previous analysis of the HIV-1 CA N128E found that this mutant reduced infectivity 50-fold compared to WT [37].

### Mutations disrupt in vitro HIV-2 CA-NC assembly

*In vitro* protein assembly assays have been very useful for analyzing protein assembly due to this being an efficient system to perturb protein-protein interactions in the absence of other cellular effects [46]. To assess the impact of selected HIV-2 CA mutations on protein interactions, a HIV-2 CA-NC construct was generated to express and purify recombinant protein for *in vitro* assembly assays. HIV-2 CA-NC WT and selected point mutants were dialyzed against assembly buffer conditions to induce helical assembly formation, and quantified by turbidity measurements at A340 nm. At this wavelength the turbidity measurement reflects the state of protein assembly from a soluble protein state to higher order oligomers [47, 48]. Based upon the severe reduction of immature particle production (**Fig. 3, 4A**), L19A, A42D, I152A, G38M and N127E were selected and the mutant protein’s ability to form helical assemblies was analyzed by taking a reading every 20 min for 2h (**Fig. 6**).

**Figure 6.**
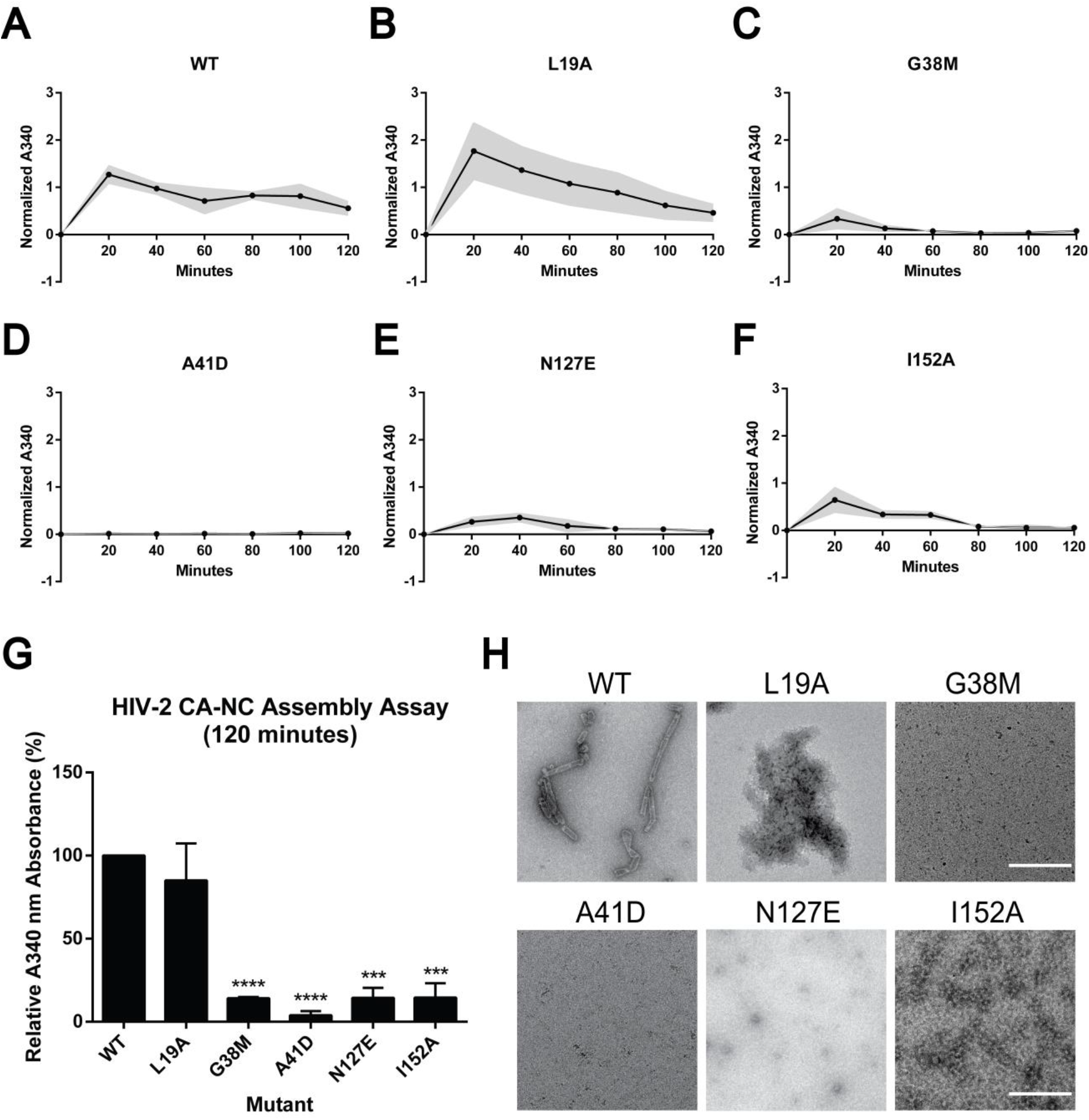
Cell-free assembly of HIV-2 CA-NC protein multimers. (A-F) Assembly kinetics of HIV-2 WT and the indicated mutants (i.e., L19A, G38M, A41D, N127E, and I152A) were monitored by measuring light scattering at 340 nm while dialyzing the reaction from the sizing buffer into the assembly buffer at the indicated times. Readings were taken every 20 min for 2 h and corrected for any changes in reaction volume due to dialysis. The grey area represents average ± standard error of the mean from three independent experiments. (G) Relative A340 nm absorbance of each HIV-2 CA mutant to WT, sampled at 120 min. Error bars represent the standard error of the mean from three independent experiments. Significance relative to WT was determined by using the unpaired t test. ****, *P* < 0.0001; ***, *P* < 0.001. (H) Negative staining of representative electron microscopy images of HIV-2 CA-NC assembly products, sampled at 120 min. Scale bar, 500 nm.

The normalized A340 nm reading for HIV-2 WT peaked at 20 min (i.e., 1.27), then slowly decreased and remained stable (i.e., 0.56) (**Fig. 6A**). HIV-2 L19A, G38M, and I152A had similar trends with WT, but with normalized A340 nm values 1.77, 0.34, and 0.65, respectively (**Fig. 6**). N127E reached a peak at 40 min, with an A340 nm value of 0.36, while A41D had low turbidity readings (i.e., < 0.36) during the experimental time course (**Fig. 6**). G38M, A41D, N127E and I152A had decreased assembly kinetics relative to that of WT.

At the final endpoint (120 minutes), G38M, A41D, N127E, and I152A had significant reductions in turbidity relative to WT, which were 7.0-fold (i.e., G38M, N127E, I152A) or 25-fold (A41D) lower than that of WT (**Fig. 6G**). These observations suggest that these mutants form unstable assembly products that disrupt CA-NC helical assembly, except for L19A. These observations also indicate that these residues play critical roles in the formation of CA-NC helical assemblies. Previous studies of HIV-1 A42D CA-NC protein assembly were also shown to impair helical assembly formation [46], which is in good agreement with our observations with HIV-2 A42D.

Visualization of these CA-NC helical assemblies by negative stain TEM reveals that HIV-2 WT CA-NC assembled into stable, regularly ordered uniform tubular structures, while G38M, A41D, N127E, and I152A did not (**Fig. 6H**). The L19A CA-NC mutant resulted in protein aggregations without uniform tubular structures, suggesting that the observed turbidity was due to large protein aggregates versus ordered helical tubes. Taken together, these results provide evidence for HIV-2 CA mutants had defects in protein interactions that prevented helical tube formation.

## Discussion

Particle morphologies among retroviral genera are quite distinct, indicating a diversity of morphologies that are different from that of HIV-1[39]. Intriguingly, HIV-2 also produces immature particles that are morphologically distinct from that of HIV-1 – *i.e.,* they possess a nearly complete immature Gag lattice, have a larger average particle diameter, and have a relatively higher average copy number of incorporated Gag [39]. Therefore, we sought out to generate a comprehensive analysis of HIV-2 Gag with a panel of HIV-2 CA mutants to analyze Gag interactions in virus particle assembly.

The primary goals of our experiments were two-fold: first, to investigate the critical role of the HIV-2 CA in Gag interactions and virus particle assembly; and second, to identify differences between HIV-1 and HIV-2 CA domains that alter particle assembly. We created a panel of 31 site-directed mutants to achieve these goals (**Fig. 1**). This panel of 31 mutants encompassed 21 alanine-scanning mutants at highly conserved residues in HIV-2 CA, and 10 site-directed mutants at non-conserved residues (*i.e.,* swapping the native HIV-2 residue to the corresponding residue observed in HIV-1 CA).

This approach allowed for an interrogation of key selected residues in the HIV-2 CA domain of Gag, based upon their predicted location at the CA intra-hexamer interface, the inter-hexamer interface, and the linker domain. To aid in our selection of mutants, specific residues were chosen based upon a structural comparison between HIV-1 and HIV-2 CA [30, 36, 37, 46, 49]. Mutations that resulted in a disruption of productive Gag interactions in immature lattice (*i.e.,* a two-fold or greater decrease in particle production) were selected for subsequent analyses. The majority of the mutants analyzed had no overall effect on immature particle production. However, seven of the mutants at conserved amino acid residues (*i.e.,* L19A, A41D, I152A, K153A, K157A, N194A, D196A) (**Fig. 2**) and two mutants at non-conserved residues (*i.e.,* G38M, N127E) (**Fig. 3A**) had a significant impact on Gag interactions and efficient particle assembly.

For mutants at the studied conserved residues and for the mutants at the HIV-2 CANTD-CACTD linker domain, there was no significant reduction in immature particle production unless they were at the MHR domain (**Fig. 2A**). Mutants at the intra-hexamer interface were observed to reduce immature particle production (**Fig. 2B**). Furthermore, mutants at the three-fold interface, but not at the two-fold interface, significantly reduced immature particle production (**Fig. 2C**), indicating that the residues at the intra-hexamer interface and at the three-fold inter-hexamer interface are critical for immature particle production and CA interactions in the Gag lattice. Importantly, immature particle production for several key mutants, including L19A, I152A and N194A, had not been previously reported.

A particularly notable discovery was two key mutants (G38M and N127E) at non-conserved residues. Intriguingly, the key features of the HIV-2 CA G38M and N127E Gag variants included a significant reduction of immature and mature particle production (**Fig. 3A, C**). This observation indicates that these mutations impact one or more steps in the particle assembly pathway. Furthermore, these mutants demonstrate a complete loss of particle infectivity (**Fig. 3D**). Gag subcellular distribution revealed an increase in nuclear and perinuclear localization for the G38M and N127E mutants relative to that of WT (**Fig. 4**). Direct visualization of the mutant immature particles by cryo-EM revealed decreased Gag lattice organization below the viral membrane and increased particle diameter for the two mutants as compared to WT (**Fig. 5**). The ability of CA to form oligomers in a cell-free assay demonstrated that these two CA mutations significantly disrupt HIV-2 CA-NC protein assembly, given that no ordered assemblies were observed (**Fig. 6**). To assess the structural basis of these results, we analyzed the molecular interfaces in HIV-1 lattice structures from both immature and mature virus particles. The HIV-1 E128 residue on helix 7 is within interaction distance (< 4Å) to E45 on helix 2 in the immature CA lattice three-fold interface. Using this structural comparison, we chose to further probe the homologous HIV-2 residue E44 and its potential interaction partner (*i.e.,* N127). Two mutations in HIV-2, E44N and N127E, resulted in significant particle reduction individually, but the double mutant (*i.e.,* E44N/N127E) had a partial rescue in particle production, suggesting that these residues have binding affinity in the immature HIV-2 Gag lattice (**Fig. 3B**). Confirmation with direct protein binding assays probing the three-fold Gag interface or solving the HIV-2 Gag immature lattice structure to near-atomic resolution would be appropriate follow-ups for explaining why these mutations altered biophysical protein binding properties.

Our results demonstrate that relatively minor changes in CA residues could profoundly impact overall particle assembly. Non-conserved residues such as G38 and N127 likely play a critically important role in HIV-2 CA structure and binding affinity. Intriguingly, the G38 and N127 residues are critical residues in the HIV-2 CA three-fold inter-hexamer interface, implying that the three-fold Gag lattice interface likely plays a critical role in HIV-2 immature particle assembly.

One potential limitation of our study is that not all residues in the HIV-2 CA domain of Gag were analyzed. Residues were identified for site-directed mutagenesis based on a variety of parameters including mutants (i) at the CA: inter-domain linker, intra-hexamer interface, and inter-hexamer interface (two-fold interface & three-fold interface); (ii) at conserved and non-conserved residues; (iii) at residues in helices and loops; (iv) and previously reported and not reported for HIV-1 CA mutagenesis studies. While not exhaustive, the panel of mutants has revealed novel residues not previously reported as important for HIV particle assembly in literature. Future studies may likely identify other important residues. Finally, analysis of novel mutants and their impact on viral infectivity through the identification of second-site reversion mutations would further aid in gaining a deeper understanding of the intermolecular interactions of Gag for virus particle assemble and infectivity.

The observations in this current study provide an important complement to structural analyses of HIV-2 immature and mature particle morphologies, and help provide important insights into the structure-function relationship between HIV particle morphology and virion infectivity. Furthermore, these studies highlight the importance of comparative virology in providing important insight into differences that can advance knowledge in our fundamental understanding of HIV assembly and replication.

## MATERIALS AND METHODS

### Plasmids, cell lines, and reagents

**The** HIV-2 Gag expression plasmids, pN3-Gag and pEYFP-N3-Gag, as well as the HIV-2 Env expression plasmid have been previously described [39]. HeLa and HEK293T/17 cells lines were purchased from ATCC (Manassas, VA) and cultured in Dulbecco’s modified Eagle medium (DMEM) supplemented with 10% fetal clone III (FC3; GE Healthcare Lifesciences, UT) and 1% Penicillin-Streptomycin (Pen Strep; Invitrogen, CA) at 37 °C in 5% CO_2_. The HIV-1 and HIV-2 single-cycle vectors, *i.e.,* pNL4-3 MIG [50] and pROD-MIG [51] have been previously described, respectively. Vector viruses were pseudotyped with the VSV-G expression construct, pHCMV-G (J. Burns, UCSD, San Diego, CA). U373-MAGI-CXCR4_CEM_ cells (NIH AIDS Reagent Program, NIAID, NIH) were maintained similarly to HEK293T/17 cells, supplemented with 1.0 µg/mL puromycin, 0.1 mg/mL hygromycin B, and 0.2 mg/mL neomycin to the medium. All cells used in this study were certified as being mycoplasma-free.

### Site-directed mutagenesis of Gag expression plasmids

A panel of 31 alanine-scanning mutants (**Fig. 1A**) were created in the HIV-2 CA domain of Gag. Among this panel, 21 alanine-scanning HIV-2 CA mutants at conserved residues were in the pN3-Gag plasmid of HIV-2 Gag (except A41D) by using the Gibson assembly method as previously described [52]. A panel of 10 non-conserved HIV-2 CA mutants were generated by changing the codons in HIV-2 to HIV-1 NL4-3 residues in the HIV-2 Gag pN3-Gag expression plasmid. Selected mutants of interest were also introduced into the pEYFP-N3-Gag, pROD-MIG, and pET28a plasmids for cellular localization, infectivity, and in vitro assembly studies, respectively. All mutants were confirmed by Sanger sequencing.

### Virus particle production

The efficiency of immature particle production was analyzed by quantifying Gag proteins in released particles by harvesting cell culture supernatants by using immunoblot analysis with a mouse monoclonal anti-HIV-1 p24 antibody (Catalogue #: sc-69728; Santa Cruz Biotechnology, TX). Briefly, the pN3-HIV-2-Gag plasmid and the HIV-2 Env expression plasmid were co-transfected into HEK293T/17 cells by using GenJet, ver II (SignaGen, Gaithersburg, MD) at a 10:1 ratio, respectively.

After 48-h post-transfection, the viral supernatants were harvested, clarified by centrifugation (1,800 × g for 10 min), and filtered through 0.2 µm filters. Next, the filtered cell culture supernatants were concentrated by ultracentrifugation in a 50.2 Ti rotor (Beckman Coulter, CA) at 211,400 × g for 90 min through an 8% Opti-prep (Sigma-Aldrich, MO) cushion. Particle pellets were resuspended in 1× STE buffer (100 mM NaCl, 10 mM Tris pH 8.0, and 1 mM EDTA) (G-Biosciences, MO). The 293T/17 cells were collected and lysed with RIPA lysis buffer and clarified via centrifugation (1800 x g for 10 min). The protein concentrations were measured by using the BCA assay (Pierce, Rockford, IL) before the samples were subjected to SDS-PAGE and then transferred to nitrocellulose membranes. The Gag proteins were detected with a 1:1,500 dilution of anti-HIV p24 antibody in 5% milk TBST (Tris-buffered saline plus Tween-20). Glyceraldehyde 3-phosphate dehydrogenase (GAPDH) was detected by using a 1:1,000 anti-GAPDH hFAB™ Rhodamine antibody (Bio-Rad, Hercules, CA) in 5% milk TBST (Tris buffer saline plus Tween-20). Membranes were washed before incubation with a goat anti-mouse StarBright™ Blue 700 secondary. Gag expression levels in cells were normalized relative to GAPDH levels, and the mutant Gag expression levels were determined relative to that of WT HIV-2 Gag.

The efficiency of mature particle production was analyzed by quantifying Gag by detection of the CA (p24) band from cell culture supernatants. Briefly, the HIV-2 pROD-MIG and the VSV-G expression plasmids were co-transfected into HEK293T/17 cells by using GenJet ver II at a 3:1 molar ratio to produce both WT and mutant virus particles. Forty-eight hours post-transfection, the cell culture supernatants were harvested, clarified by centrifugation (1,800 × g for 10 min), and filtered through 0.2 µm filters. The supernatants were then concentrated by ultracentrifugation in a 50.2 Ti rotor (Beckman Coulter, CA) at 211,400 × g for 90 min through an 8% Opti-prep (Sigma-Aldrich, St. Louis, MO) cushion. Capsid proteins were detected with a 1:1,500 anti-HIV p24 antibody. Glyceraldehyde 3-phosphate dehydrogenase (GAPDH) was detected with 1:1,000 anti-GAPDH hFAB™ Rhodamine antibody (Bio-Rad). The CA levels from cells were normalized relative to GAPDH levels, and the CA mutant expression levels were determined relative to that of WT HIV-2 Gag. Membranes were washed before incubation with 1:2,500 goat anti-mouse StarBright™ Blue 700 (Bio-Rad).

The membranes from immunoblot analyses were imaged by using a ChemiDoc Touch system (Bio-Rad, CA) and analyzed with ImageJ. Results were analyzed by using GraphPad Prism 6.0 (GraphPad Software, Inc., CA). Relative significance between a mutant and WT was determined by using an unpaired t-test. Immunoblot analyses were done by conducting three independent replicates.

### Gag subcellular distribution analysis

Gag subcellular distribution was analyzed by quantifying the degree of Gag puncta formation in cells by using confocal laser scanning microscopy to localize Gag-eYFP as previously described [53]. Briefly, HeLa cells were cultured in six-well plates on 1.5 standard glass coverslips coated with poly-L-lysine as previously described [53]. HeLa cells were transiently transfected with EYFP-tagged Gag; untagged Gag expression plasmids at a 1:3 molar ratio by using GenJet, ver II (SignaGen, Gaithersburg, MD). At 48-h post-transfection, cells were stained by using DAPI (Thermo Fisher Scientific, MA) and ActinRed 555 (Invitrogen, CA) prior to before fixation with 4% paraformaldehyde (Thermo Fisher Scientific, MA). Cells were imaged by using a Zeiss LSM 700 confocal laser scanning microscope with a Plan-Apochromat 63×/1.40-numeric-aperture (NA) oil objective at 1.2× zoom (Carl Zeiss, Oberkochen, Germany). At least 5 individual cells were imaged from three independent experimental replicates, for a total of 15 cells for each Gag mutant and WT Gag.

### Cryo-EM analysis of particle morphology

The pN3-HIV-2-Gag plasmid and the HIV-2 Env expression plasmids were co-transfected into HEK293T/17 cells by using GenJet, ver II at a 10:1 molar ratio as previously described [39]. At 48-h post-transfection, cell culture supernatants were harvested and centrifuged at 1,800 × g for 5 min and followed by passing through a 0.2 µm filter. Particles were then concentrated by ultracentrifugation in a 50.2 Ti rotor at 211,400 × g for 90 min through an 8% Opti-prep cushion. Particle pellets were resuspended in about 200 µl of STE buffer prior to ultracentrifugation through a 10% to 30% Opti-Prep step gradient at 301,090 x g for 3 h. The particle band was removed from the gradient by puncturing the side of the centrifuge tube with a hypodermic syringe needle and pelleted in STE buffer at 267,636 × g for 1 h by using an SW55 Ti rotor. The resulting particle pellet was then resuspended in approximately 10 µl STE buffer and frozen at -80°C until analyzed by cyro-EM. Experiments were done in triplicate. Samples were initially screened by negative staining TEM with 0.75 % (w/v) uranyl formate by using a 120 kV Tecnai Spirit TEM [54].

Cryo-EM analysis of particles were done as previously described [39, 53, 55]. Briefly, particle samples were thawed on ice, and approximately 3.5 µL of purified virus particles were applied to freshly glow-discharged (10 mA for 30 sec using a Leica Ace600 glow-discharger) Quantifoil R2/1 300-mesh holey carbon-coated copper grids. The grids were then blotted for 4-10 sec with filter paper at 19°C with 85% relative humidity and plunge-frozen in liquid ethane using a FEI Mark III Vitrobot or Leica GP-2 grid plunger. The frozen grids were stored in liquid nitrogen until imaging analysis.

Cryo-EM analysis was done by using a Tecnai FEI G2 F30 FEG transmission electron microscope (FEI, Hillsboro, OR) at liquid nitrogen temperature operating at 300 kV. Images were recorded at a nominal magnification of 39,000x and 59,000x magnification under ∼25 electrons/Å^2^ conditions at 1 to 5 μm under-focus by using a Gatan Slow Scan 4k by 4k charge-coupled-device (CCD) camera or a Gatan K2 Summit direct electron detector (Gatan Inc., Pleasanton, CA). At least 50 individual immature particles of each mutant were measured using ImageJ software. Two perpendicular diameters were measured and averaged for each particle. Particle morphologies were qualitatively characterized.

### Virus infectivity assay

The HIV-2 pROD-MIG and VSV-G expression plasmids were co-transfected into HEK293T/17 cells by using GenJet, ver II at a 3:1 molar ratio to produce WT and mutant particles. Forty-eight hours post-transfection, cell culture supernatants were harvested, clarified by centrifugation (1,800 × g for 10 min), and filtered through a 0.2 µm filter. U373-MAGI-CXCR4 cells were plated in a 12-well plate and 1 ml of concentrated cell culture supernatant along with 1 ml fresh medium were added per well. Each mutant was analyzed in a quadruplicate. The cells were collected to detect virus-infected cells via flow cytometry by using a BD LSR II flow cytometer (BD Biosciences, San Jose, CA)) at 48-h post-infection [45]. Data were analyzed by using FlowJo v.7 (FloJo Inc, Ashland, OR). Infected cells were determined by identification of all quadrants expressing either or both fluorescence markers (i.e., mCherry, GFP, and both mCherry/GFP) relative to that of WT vector virus. Relative infectivity was normalized to particle production as determined by immunoblot analysis of cell culture supernatants. Experiments were conducted in triplicate

### Cell-free HIV-2 CA-NC assembly analysis

**The** HIV-2 CA-NC (Gag amino acids 136-431) was cloned into the pET28a bacterial expression plasmid, with mutants created by using the Gibson assembly procedure. Protein is expressed in *E. coli* by using as previously described [47, 48]. Briefly, the CA-NC protein was expressed in BL21 (DE3) RIP pLysS *E. coli* grown in 200 mL of ZY-auto induction media in a platform shakers for 16 h at 37C [56]. Cells were pelleted and resuspended in 50 mL lysis buffer per 200 mL culture (500 mM NaCl, 25 mM Tris pH 7.5, 1 µM ZnCl2, 10 mM β-Mercaptoethanol (BME)) and then exposed to flash freezing to aid in cell lysis. After thawing, cells were lysed by adding 10 mg lysozyme, 0.1% Triton (v/v), and sonicated for 10 sec on/off cycles at 40% amplitude for three, 5 min treatments to reduce viscosity. The cell lysate was clarified by centrifugation 12,000 × g 40 min. Nucleic acids were removed by using 0.11 (v/v) 2M ammonium sulfate and 0.11 (v/v) polyethyleneimine (PEI), and protein subsequently precipitated by using saturated ammonium sulfate with 1/3 (v/v) of the total supernatant volume. The precipitated protein was pelleted by centrifugation at 7,500 × g spin for 15 min. The precipitate was then resuspended in 50 mM NaCl, 25 mM Tris, 5 µM ZnCl2, 10 mM BME, pH 7.5, and further dialyzed against 1L for 1 h at 4C. The protein was concentrated using an Amicon ultra-15 centrifugal filter unit concentrator (Millipore, Burlington, MA) and loaded onto an ion-exchange column (HiTrap SP FF, Cytiva, Global Life Sciences Solutions USA LLC, Marlborough, MA). Protein was eluted by using a linear NaCl gradient from 50-700 mM. Fractions containing the protein were pooled, concentrated to ∼10 mg/mL, and flash-frozen in liquid nitrogen. Only samples below a 0.6 A260/A280 ratio were used for further analysis to ensure no significant nucleic acid contamination was affecting the assembly kinetics or products.

The CA-NC assembly reactions were done using purified protein diluted to 50 µM in 250 mM NaCl, 50 mM Tris, pH 8. A 50-mer oligonucleotide consisting of 25 (GT) repeats was added at a final concentration of 5 µM. The protein was then dialyzed against assembly buffer (2M NaCl, 50 mM Tris, 150 µM IP6, 5 µM ZnCl2, pH 8) at room temperature in Pierce^TM^ Slide A Lyzer^®^ Mini Dialysis Units (Thermo Fisher Scientific, Waltham, MA). Turbidity measurements at A340 nm were collected in order to quantify the reaction progress by using NanoDrop 1000 Spectrophotometer (Thermo Fisher Scientific, Waltham, MA). Spectrometry readings were collected every 20 min for 2 h, and corrected for any changes in reaction volume due to dialysis.

After 2 hours of dialysis, 3 µl of each reaction were spotted onto EMS CF300-CU grids (Ted Pella, Redding, CA) for 2 min. Samples were then blotted with filter paper, washed 3x in deionized water, blotted, and stained in 0.75% (w/v) uranyl formate for 2 min. Samples were imaged by using a FEI Technai Spirit Bio-Twin transmission electron microscope at 120 kV.

## Acknowledgments

This research was supported by NIH grant R01 AI150468 (to L.M.M.). N.T. was supported by NIH grants T32 DA007097, F32 AI150351, and American Cancer Society Postdoctoral Fellowship PF-21-189-01-MPC. W.G.A. was supported by NIH grant NIH T32 AI083196. TEM was carried out in the Characterization Facility, University of Minnesota, which receives partial support from NSF through the MRSEC and NNCI program.

---

**Figure S1.**
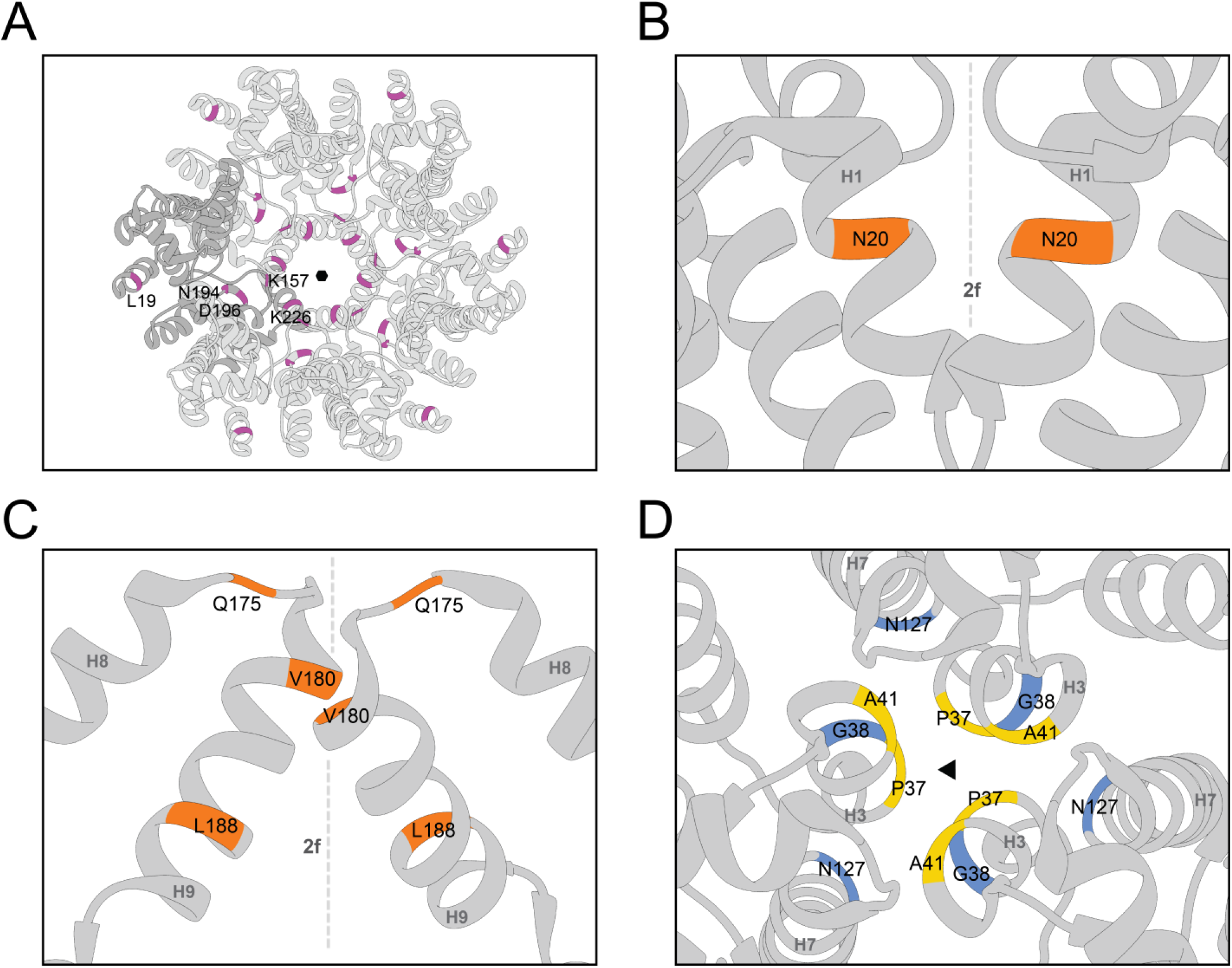
Mutagenesis sites of the CA hexamer and inter-hexamer interface (related to Fig.1). Mutagenesis at intra-hexamer interface (magenta), two-fold interface (orange), three-fold interface (blue), and non-conserved residues (yellow). (A) Immature HIV-1 CA hexamer with 5 mutants at the intra-hexamer interfaces. (B) CANTD two-fold interface (Helix 2). (C) CACTD two-fold interface (Helix 9). (D) CA three-fold interface. PDB ID: 5L93.

**Figure S2.**
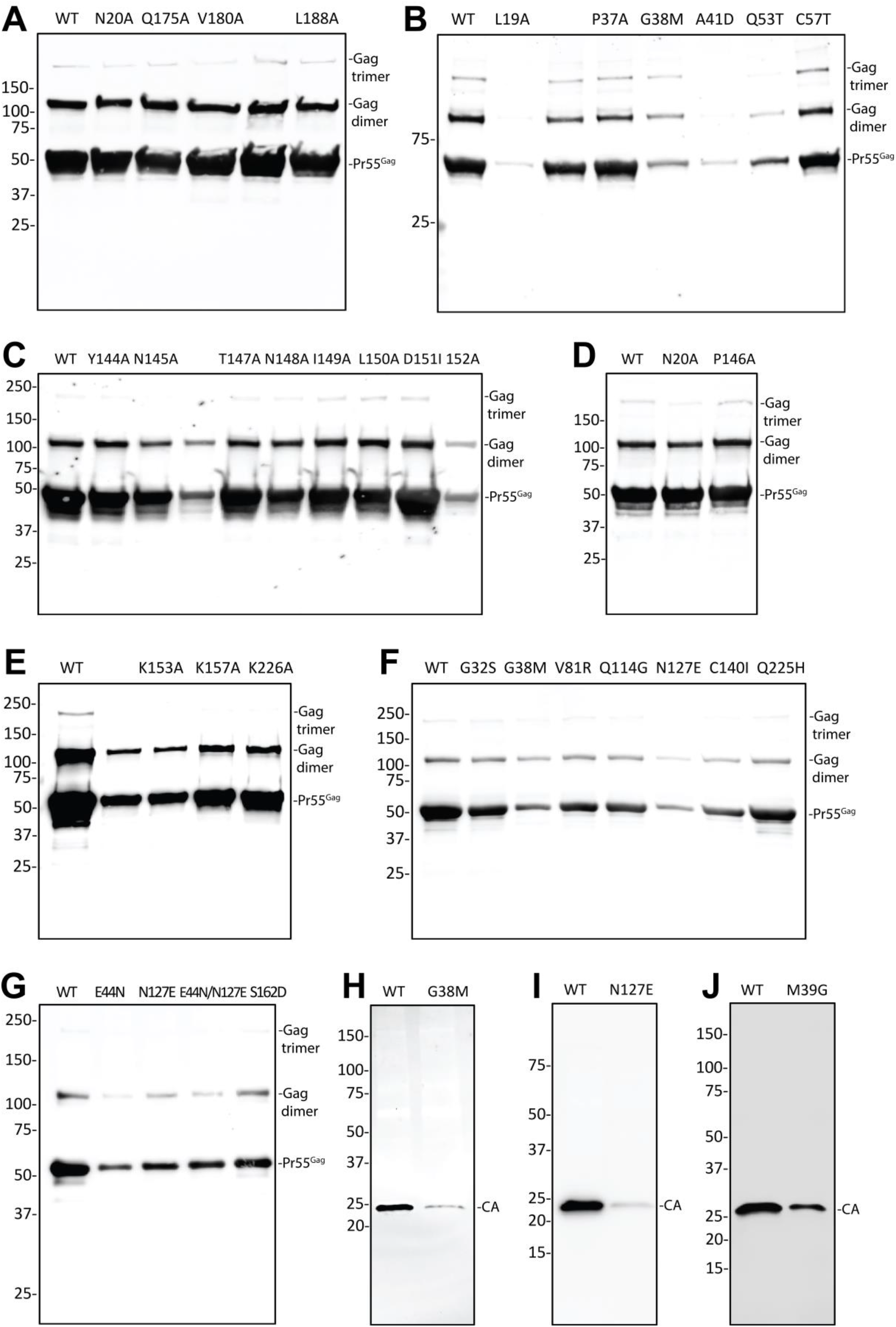
Representative immunoblot images from three independent experiments (related to Fig. 2 and Fig. 3). (A-G) Immature particle production for HIV-2 CA mutants at conserved and non-conserved amino acid residues. The efficiency of immature particle production was analyzed by quantifying Gag proteins with a monoclonal anti-HIV p24 antibody. (H-J) Mature particle production for HIV CA mutants. The efficiency of mature particle production was analyzed by quantifying Gag by detection of the CA (p24) band from cell culture supernatants.

